# Deep mapping of the *endomembrane system* of cerebellar Purkinje neurons

**DOI:** 10.1101/2025.07.08.663701

**Authors:** Matthias G Haberl, Silvia Viana da Silva, Sebastien Phan, Eric Bushong, Thomas Deerinck, Mark H Ellisman

## Abstract

Neuronal function relies on the precise spatial organization of intracellular membrane-bounded organelles involved in anabolism and Ca^2+^ sequestration, such as the Golgi apparatus, mitochondria and the endoplasmic reticulum (ER), along with structures involved in catabolism, such as lysosomes. Despite their known roles in energy supply, calcium homeostasis, and proteostasis, our understanding of how the anabolism-linked organelles are structurally arranged within neurons remains incomplete. Due to the tremendous complexity in the morphologies and fine structural features and interwoven nature of these intracellular organelles, particularly the ER, our understanding of their structural organization is limited, particularly, with regard to quantitative assessments of their sites of interaction and accurate measures of their volumetric proportions inside of a single large neuron. To approach this challenge, we used serial block-face scanning electron microscopy (SBEM) to generate large-scale 3D EM volumes and electron tomography on high-pressure frozen tissue of the rodent cerebellum, including the largest cells in the vertebrate brain, the cerebellar Purkinje neuron as well as the most abundant cell type in the vertebrate brain, the much smaller cerebellar granule neuron. We reconstructed the neuronal ultrastructure of these different cell types, focusing on the ER, mitochondria and membrane contact sites, to then characterize intracellular motifs and organization principles in detail, providing a first full map to quantitatively describe a neuronal *endoarchitectome*. At the gross level organization, we found that the intracellular composite of organelles are cell type specific features, with specific differences between Purkinje neurons and Granule cells. At the level of fine structure, we mapped ultrastructural domains within Purkinje neurons where ER and mitochondria associate directly. In addition to cell type specific differences, we observed significant subcellular regional variation, particularly within the axon initial segment (AIS) of Purkinje neurons, where we identified ultrastructural domains with sharply contrasting distributions of ER and mitochondria. These findings suggest a finely tuned spatial organization of organelles that may underpin the distinct functional demands along the axon. We expect that our subcellular map, along with the methods developed to obtain these maps, will facilitate future studies in health, aging and disease to characterize defined features, by developing a framework for quantitative analysis of the neuronal ultrastructure.

## Introduction

The endoplasmic reticulum (ER) in neurons is a large and complex organelle, which starting from the nuclear envelope (*1*) meanders throughout soma, axons (*2*, *3*) and dendrites, and even into the presynaptic boutons (*4*) and postsynaptic spines (*5*, *6*). The ER is linked with the Golgi apparatus (*7*) and has numerous contact sites with the cell membrane and mitochondria (*8*, *9*) which are critical for its function in regulating intracellular calcium. To this end, it has a lumen rich in sequestered Ca^2+^ (*2*), which is released through its membrane by Ca^2+^ channels, specifically inositol trisphosphate receptors (IP3Rs) and ryanodine receptors (*10*). Within the soma of neurons, the ER is a major constituent in volume and surface, yet it has received much less attention compared to the much smaller, less prominent ER in spines, referred to as spine apparatus (see e.g. (*6*, *11*) and reviewed in (*12*)), since its dimensions are difficult capture and reconstruct at sufficient resolution it poses a tremendous problem for segmentation at scale. Anatomically there are a few broadly defined configurations of ER in cells and neurons, besides the nuclear envelope, namely sheets and tubules (*13*, *14*). Even though we know that also local domains can influence Ca^2+^-signals (*15*), the precise three-dimensional distribution of the ER and the occurrence of intracellular motifs in neurons from brain tissue have so far been difficult to quantify.

To address these questions about the ultrastructural organization in neurons, we acquired high-resolution multi-tilt tomograms and serial block-face scanning electron microscopic (SBEM) volumes (*16*) of cerebellar tissue. We then performed large-scale 3D reconstructions of the cellular constituents including mitochondria and the endoplasmic reticulum of granule cells and Purkinje neurons. These two cell types represent extremes in neuronal architecture, with granule cells among the smallest, and Purkinje neurons among the largest and most morphologically complex in the brain. Notably, the Purkinje neuron is a prominent inhibitory interneuron whose physiology, gene expression, and role within neural circuits have been extensively characterized across vertebrate species and throughout evolution. Based on our 3D segmentations, we performed quantifications of the intracellular architecture of Purkinje neuron somata and the axon initial segment (AIS). We then characterized in detail the intracellular ultrastructure of the neuronal soma of Purkinje neurons, which are large GABAergic inhibitory neurons, providing the sole output of motor coordination from the cerebellar cortex (*17*).

Intracellular morphological organization principles of neurons are critical to understand spatial constraints imposed on intracellular functions and intracellular signaling traveling of messenger molecules that are important for the communication in the brain. Large-scale volume electron microscopy (vEM) is rapidly becoming widely used to obtain dense connectomic reconstructions across species, brain regions and physiological states (*18–23*). Similarly whole-cell organelle segmentations of these vEM data are being carried out on cultured cells or assemblies of cells in somatic organs, like those of pancreatic islets, at larger scales (*24*–*26*), propelled by advances in deep learning, computational resource growth and AI. The work described here demonstrates the application of these strategies and lays out a foundation for future studies with examples of quantifiable parameters relevant to the intracellular organization, in particular for neuronal somata, axons and dendrites in health and disease. We expect that our work thus provides a first ultrastructural blueprint of a neuronal *endoarchitectome*, which provides detailed information required for computational spatial simulations and can contribute to a next level of cell type characterizations, across ages, during dynamics of activities and variations in underlying health.

## Results

### Large- and fine-scale reconstructions of the 3D intracellular ultrastructure of the cerebellar cell somata from volumetric electron microscopy datasets

Reconstructing the intracellular features of neuronal somata, such as the endoplasmic reticulum and mitochondria, is a multiscale problem spanning from the few nm-scale of the ER diameter to tens of µm to capture the profile of intracellular distributions. In order to generate large-scale 3D maps of the ultrastructural features of Purkinje neuron somata we used serial block-face scanning electron microscopy (SBEM) of *en-bloc* stained mouse cerebellar cortex tissue for large-scale measurements to EM volumes (**Figure 1**). We acquired two SBEM microscopy datasets: one from the cerebellum of an adult male mouse with a size of 57.17 × 76.22 × 17 μm^3^ where we segmented the cellular ultrastructure of Purkinje neurons (**Figure 1 b-e**) and granule cells (**Figure 1 f-h**) and one dataset from the cerebellum of an adult rat with a size of 189 × 118 × 60 μm^3^, which includes the axon initial segment. The subsequent 3D segmentations of the electron microscopic data volumes were performed by training deep neural networks models with CDeep3M (*27*) for segmenting different cellular structures and subsequent manual proof editing of the computational segmentations by expert annotators. In order to gain insights about cell type dependent variation of the intracellular composition, we first analyzed the volume of soma, nuclei, cytoplasm ER and mitochondria in Purkinje neurons (**Figure 1i**) and granule cells (**Figure 1j**) based on the proof-edited SBEM segmentations. When calculating the respective somatic volume fraction occupancy by ER and mitochondria, we found that even though a Purkinje neuron has ∼80 times more mitochondria (i.e. >2700 versus 34 ± 5) than a granule cell, the relative volume fraction occupied by mitochondria was similar in between Purkinje (9.4%) and granule cells (10,0 ± 1,1%) in the same imaging volume (**Figure 1k**). Whereas a comparison of ER occupancy revealed that the ER occupied 12.3% in the Purkinje, but only 5,8 ± 0,9 % in the granule cells (**Figure 1l**).

**Figure 1.**
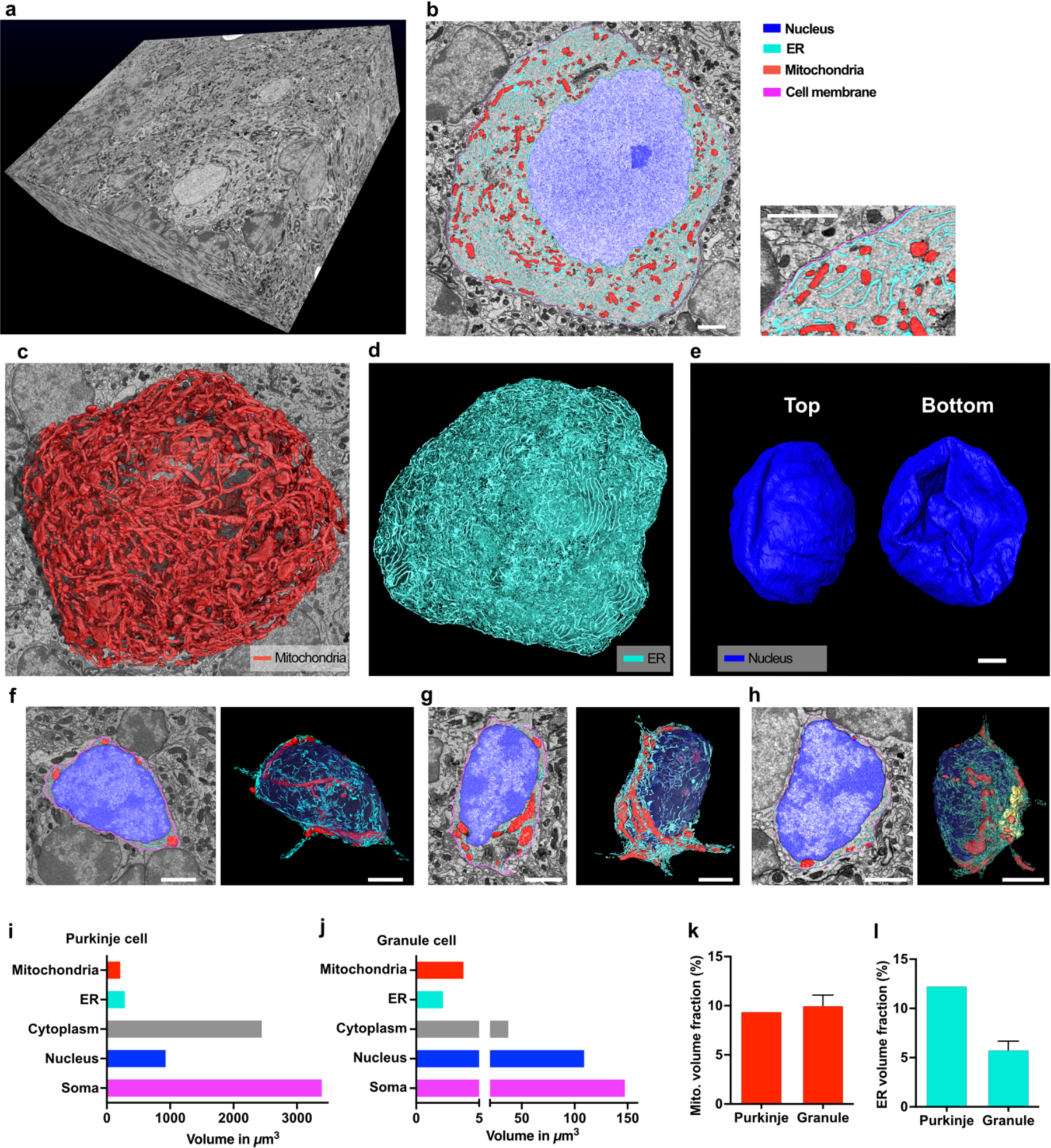
Large-scale 3D reconstructions of the intracellular ultrastructure of cerebellar Purkinje neurons and granule cells. Individual 2D view of (**a**) electron tomogram of Purkinje neuron soma acquired from high-pressure frozen freeze substituted cerebellar tissue. (**b**) Intracellular structures including the endoplasmic reticulum were computationally segmented and manually proof edited before quantifications. (**c**) Overlay in 2D of the tomogram displaying segmentations of the ER (cyan), mitochondria (red), nucleus (blue), golgi apparatus (yellow). (**d**) Overlay of 3D surface meshes of these intracellular structures and the cell membrane from one electron tomography volume. (**e**) Volume view of serial block-face scanning electron microscopy, performed on an *en-bloc* stained cerebellar tissue containing Purkinje neurons and granule cells. Example 2D planes of (**f**) Purkinje neuron soma and (**g**) granule cell soma, both overlayed with ER (cyan), mitochondria (red), nucleus (blue) and cell membrane (yellow) segmentations. 3D surface meshes of the Purkinje neuron (**h**) mitochondria (red) and (**i**) ER (cyan). Quantification of the absolute volume of the segmented organelles for the (**j**) Purkinje neuron soma and (**k**) granule cell soma. Direct comparison of the volume fractions of intracellular structures of Purkinje neurons and granule cell soma shows (**l**) similarity in mitochondria content and (**m**) a large difference in the ER content between the two cell types. Scale Bars (b-h): 2µm

In order to support our analysis of fine-scale intracellular structural features at higher resolution, we also acquired multi-tilt electron tomography datasets of high-pressure frozen, freeze substituted tissue, of cerebellar mouse cortex, a technique which can be scaled to perform large scale high resolution connectomics (*28*).

We acquired, segmented and reconstructed five three-dimensional, high-resolution, multi-tilt electron tomography volumes, at the dendritic and the axonal poles of cerebellar Purkinje neurons (**Figure 2 a-c**). The electron tomography volumes were acquired from following areas: Volume 1, 2 and 3 were acquired at the dendritic pole of Purkinje neurons in the mouse cerebellum, Volume 4 and 5 were acquired at the axonal pole of Purkinje neurons in the mouse cerebellum (**Supplementary Figure 1**).

**Figure 2.**
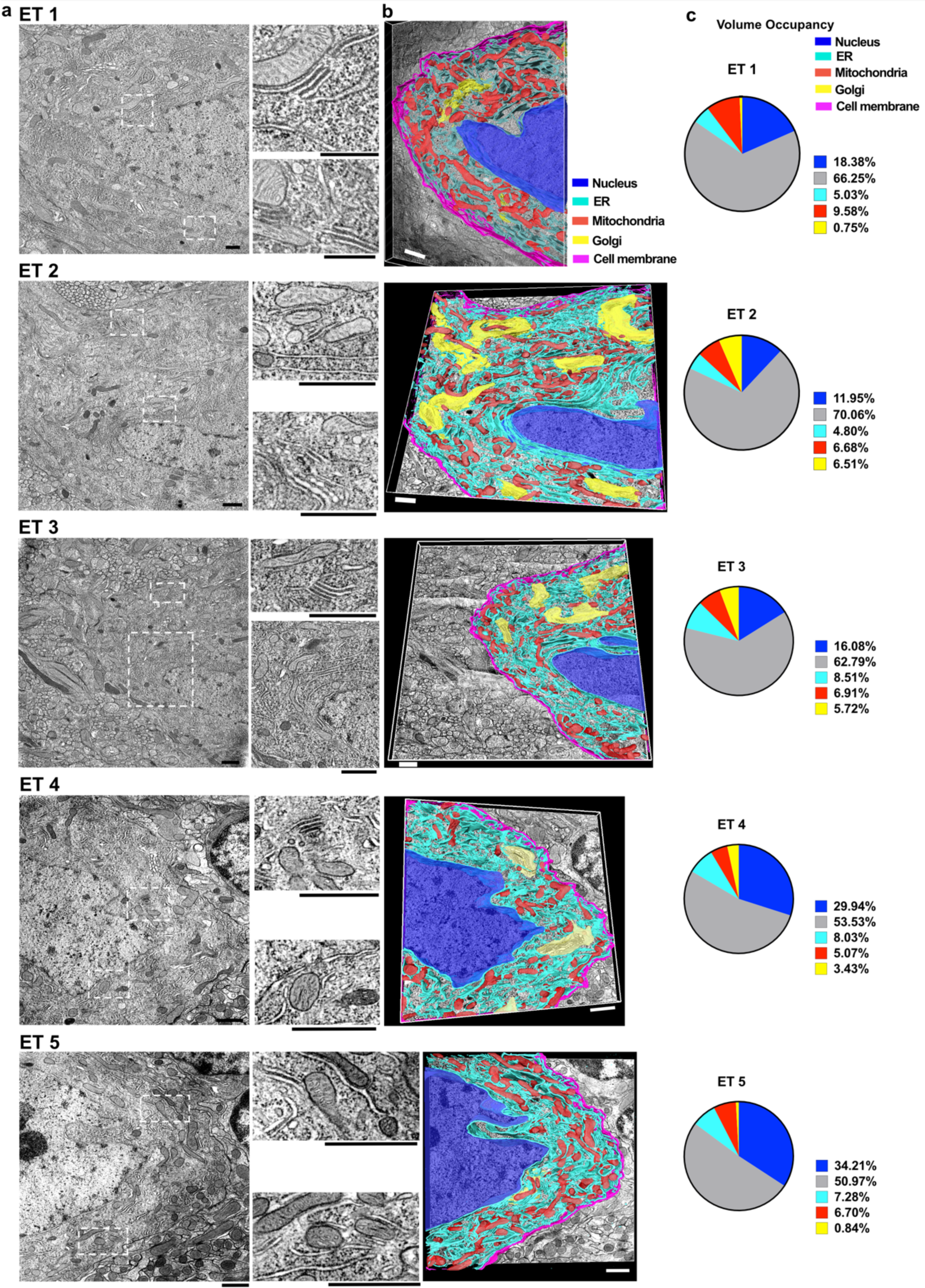
Segmentation of the intracellular architecture of Purkinje neurons based on five semi-thick electron tomography volumes. (**a**) 2D views from electron tomography volumes (ET 1-5) of Purkinje neuron soma acquired from high-pressure frozen freeze substituted cerebellar tissue. Based on these the intracellular structures including the endoplasmic reticulum were computationally segmented and manually proof edited before quantifications. (**b**) Overlay of 3D surface meshes based on segmentations of the cell membrane (magenta), ER (cyan), mitochondria (red), nucleus (blue), Golgi apparatus (yellow) showing the intracellular arrangement in the electron tomography volumes. (**c**) Volume occupancy ratios within the Purkinje neuron in each of the electron tomography volumes of intracellular organelles: nucleus, ER, mitochondria and Golgi apparatus. Cytoplasm volume was calculated as volume enclosed by the cell membrane without the volume of nucleus, ER, mitochondria and Golgi apparatus. Scale Bars: 1µm

### Somatic ER of Purkinje neurons is omnipresent and also oriented perpendicular to the plasma membrane

To characterize the morphological features of the ER, we first performed automated measurements on the reconstructions of the SBEM data where we find an average ER diameter of 44.09nm ± 12.39 nm (**Figure 3a**), which is in line with results from our high-resolution tomography data (average ER diameter of 39.25 nm ± 10.55 nm; **Supplementary Figure 2**), and an average length of in plane ER strands of 426.8 nm ± 580.5 nm (**Figure 3b**). Interestingly, we find a significant proportion of the somatic ER in the Purkinje neuron falls in the category of narrow ER (e.g. 26.9% 20-30nm in ET2; **Supplementary Figure 2**), which was suggested to be a property of axonal ER only recently (*29*).

**Figure 3.**
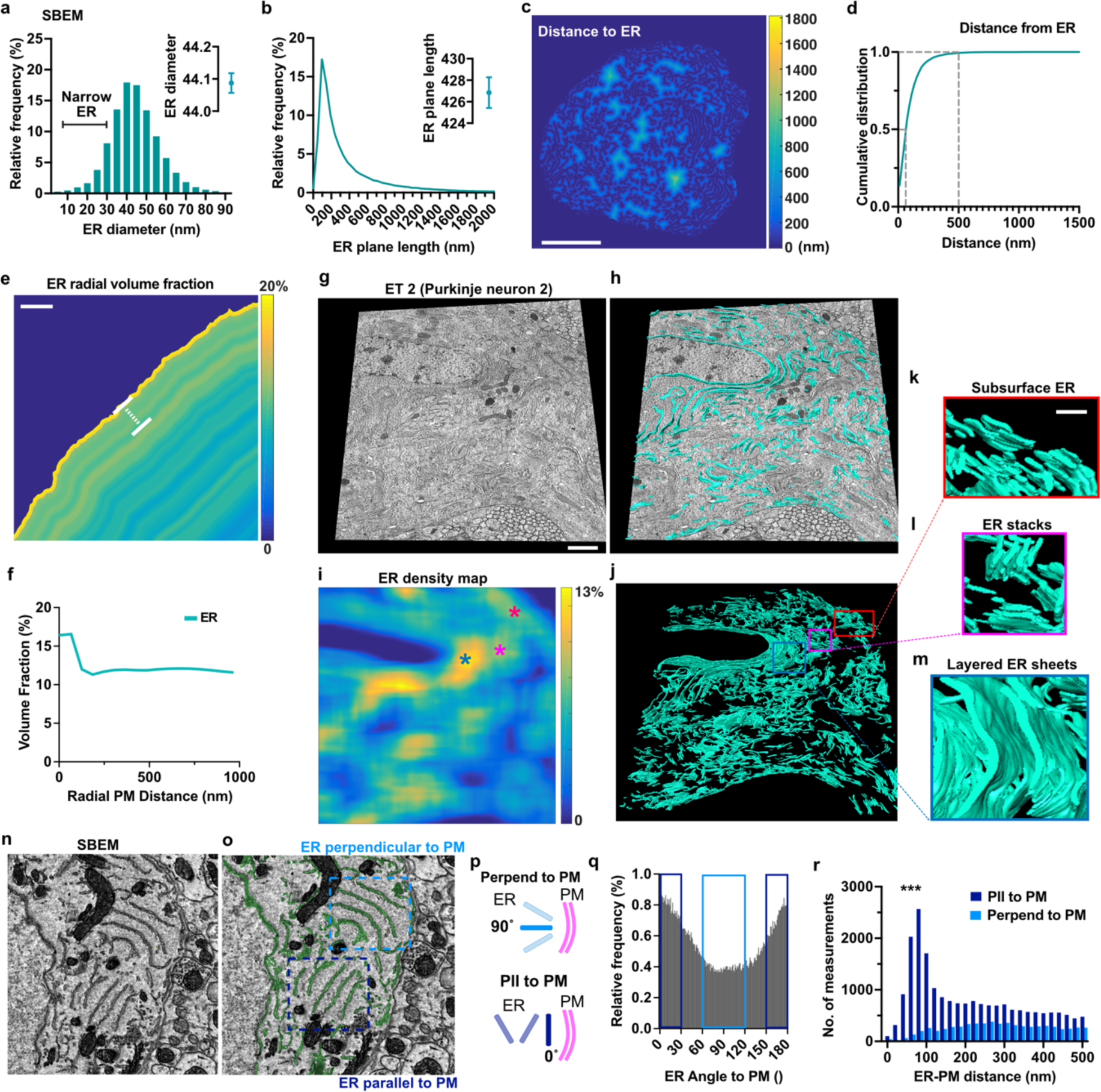
ER is also oriented perpendicular to the somatic plasma membrane of Purkinje neurons. Analysis of the ER morphology of the electron tomograms shows (**a**) ER diameter of 39.25 nm ± 10.55 nm (**b**) 426.8 nm ± 580.5 nm 2D plane ER strand length. (**c**) Heatmap plot of ER distance maps, shows if intracellular areas are void of ER (light colors). (**d**) The cumulative ER distance plot across the entire soma of the SBEM reconstruction reveals that the majority (>50%) of the Purkinje neuron soma has ER in close proximity (within 55nm) and almost all the soma (99%) has ER within 500nm. (**e**) The ER radial heat map (of a single example plane) and (**f**) the cumulative radial distribution plot (across the entire soma) display the ER volume fraction of the cell at each distance of the surface and reveal a high content of subsurface ER (within 100nm from the PM). Based on a (**g**) high-resolution electron tomography volume the (**h**) ER segmentations were transformed into (**i**) density maps which reveals that ER density peaks (light colors, yellow) are correlated to the occurrence of structural motifs of the ER, in particular subsurface ER (red label in **i-j**; and m), ER stacks (magenta label in **i-j**; l) and layered ER sheets (blue label in **i-j**; m). (**n-r**) The orientation of ER strands was quantified relative to the PM and (**p-q**) defined as parallel (pll; dark blue) or perpendicular (perpend; light blue) to the PM. (**p**) Exemplified schema of the calculation of the different angles of ER stacks. (**q**) Distribution of ER angle calculated relative to the plasma membrane (PM). (**r**) An overall preference of the ER to orient in parallel to the PM particularly at close distances to the PM is evident (ratio of pll to perpend: 93% to 7%) whereas further away from the PM the fraction of perpendicular ER increases (ratio of pll to perpend: 64% to 36%). Scale Bars (c): 5µm; (e, g-j): 1µm; (k-m): 250nm

Since the ER has several critical roles in the neuronal signaling, we wanted to determine if this implies that the structure is well distributed throughout the entire soma (excluding the interior of the nucleus), or if areas with little or no ER exist. Performing a distance analysis (heatmap plotted in **Figure 3c**), we found that indeed nearly all of the soma has ER within a distance of 250nm (93.44% of soma area has ER within 250nm and 99.37% of soma area has ER within 500nm) (**Figure 3d**). In summary, this means that ER is omnipresent throughout the soma of the Purkinje neurons and nearly all areas in the soma have ER within a short distance, underlining its critical role for cellular processes.

Since we found that the ER is omnipresent throughout the neuronal soma, we wanted to test if this also entails that it is uniformly distributed. Therefore, we performed a radial analysis of ER-occupancy (normalized by the volume at each distance). We found that the most dominant feature in this radial distribution is a prominent occupancy of ER in the space between 0-100nm near the PM (**Figure 3e**), and we encounter 65.71% volume fraction in 0 - 100nm (**Figure 3f**). This is in line with recent findings in the nucleus accumbens where the PM of the neuronal somata of neurons were even found to be covered with ER to a much larger extent than spines, dendrites, and axons (*9*). Since this type of ER in proximity to the PM - also called cortical or subsurface ER - have been shown to have a critical role for intracellular calcium signaling (*30*) and for the replenishment of ER calcium stores (*31*), we went on to measure the ratio of the PM, which is in close proximity to ER. We found that 4.79 % of the PM is covered with subsurface ER withinin 30nm distance in the electron tomogram data. This area fraction is in line with what has been described for the soma of other neuron types (for comparison: 4-12% within 30nm distance in neurons in the nucleus accumbens (*9*)), possibly reflecting the importance of ER-PM contacts for the neuron physiology in Purkinje neuron.

To further improve our understanding if somatic domains exist within the intracellular distribution of the ER, we took advantage of the high-resolution electron tomography (**Figure 3g**) and the ER reconstructions (**Figure 3h**) and performed a density analysis within the Purkinje neuron (**Figure 3i**), which revealed that peaks in the ER density maps are caused by particularly organized structural motifs of the ER (**Figure 3j**), namely layered subsurface ER (**Figure 3k**), ER stacks (**Figure 3l**) and layered ER sheets (**Figure 3m**), which is in line with known ER assemblies (*13*).

However, contrary to textbook descriptions of the ER, which often portray an exclusive parallel orientation to the nuclear envelope and cell membrane, we found in all our EM datasets that this parallel orientation is highly intermingled with ER oriented in any other orientation angle towards the PM (**Figure 3n-o**). To determine if these angles are distributed completely at random or if a preferential ER organization exists, we went on to measure the orientation angle of ER strands in the mouse SBEM volume in relation to the plasma membrane as well as in relation to the nuclear envelope. We define the parallel orientation to the plasma membrane as 0° whereas a perpendicular orientation as 90° (**Figure 3p**). While we find a preference towards orientation angles that are nearly parallel to the plasma membrane and nuclear membrane (45.1% of ER strands are within 0°± 30° angle difference between ER and PM), we also find that a large proportion of ER is oriented perpendicular (24.3% within 90°± 30° angle difference between ER and PM) or at any angle between these two extremes (**Figure 3q**). This suggests that outside of the parallel orientation, there is no clear preference for another orientation angle and all configurations occur. Importantly however, we found that the relative frequency at which the preferred orientation (parallel to the PM) occurs is dependent on the PM-ER distance and that the relative frequency of ER perpendicular to the PM increases at larger PM-ER distance (**Figure 3t**), making consistently above 30% at >200nm (mean: 33% between 200-500nm) distance from the PM.

## Mitochondria ultrastructure in the Soma of Purkinje neurons

Since the exact distribution and morphologies of mitochondria have important implications for the energy supply of different intracellular domains, and have not been described for Purkinje neurons, we next aimed to perform a detailed characterization of this cellular ultrastructure based on the 3D reconstructions of 2712 mitochondria (**Figure 4a**), with a density of 0.79 mitochondria / µm^3^ in the Purkinje neuron soma. Across the mitochondria we found an average diameter of 132 nm ± 1.16 nm (**Figure 4b**) and length of 2533nm ± 566nm (**Figure 4c**) and an average volume of 0.11 µm^3^ ± 0.02 µm^3^ (**Figure 4d**). Since the Mitochondria Complexity Index (MCI) has been found to vary and was implied in ageing, we wanted to test if the MCI is similar across cell types and if it is therefore comparable in Purkinje neurons. The average MCI in our Purkinje neurons was 5.34 ± 0.94 (**Figure 4e**), which is similar to the recently described MCI in hippocampal CA1 and DG cells (*32*). Based on our findings we developed an average shape descriptor (with standard deviation) of somatic mitochondria in the Purkinje cell bodies (**Table 1**). We further determined the orientation of mitochondria in the tomography volumes and in the SBEM dataset, and found that unlike the ER, that mitochondria have no clear preference to orientate parallel relative to the cell membrane (**Figure 4g-i**). We thus expect that the orientation of the ER may influence the neuron function, not only at the level of the diffusion of signaling molecules but potentially also more directly at the level of Calcium signaling.

**Figure 4.**
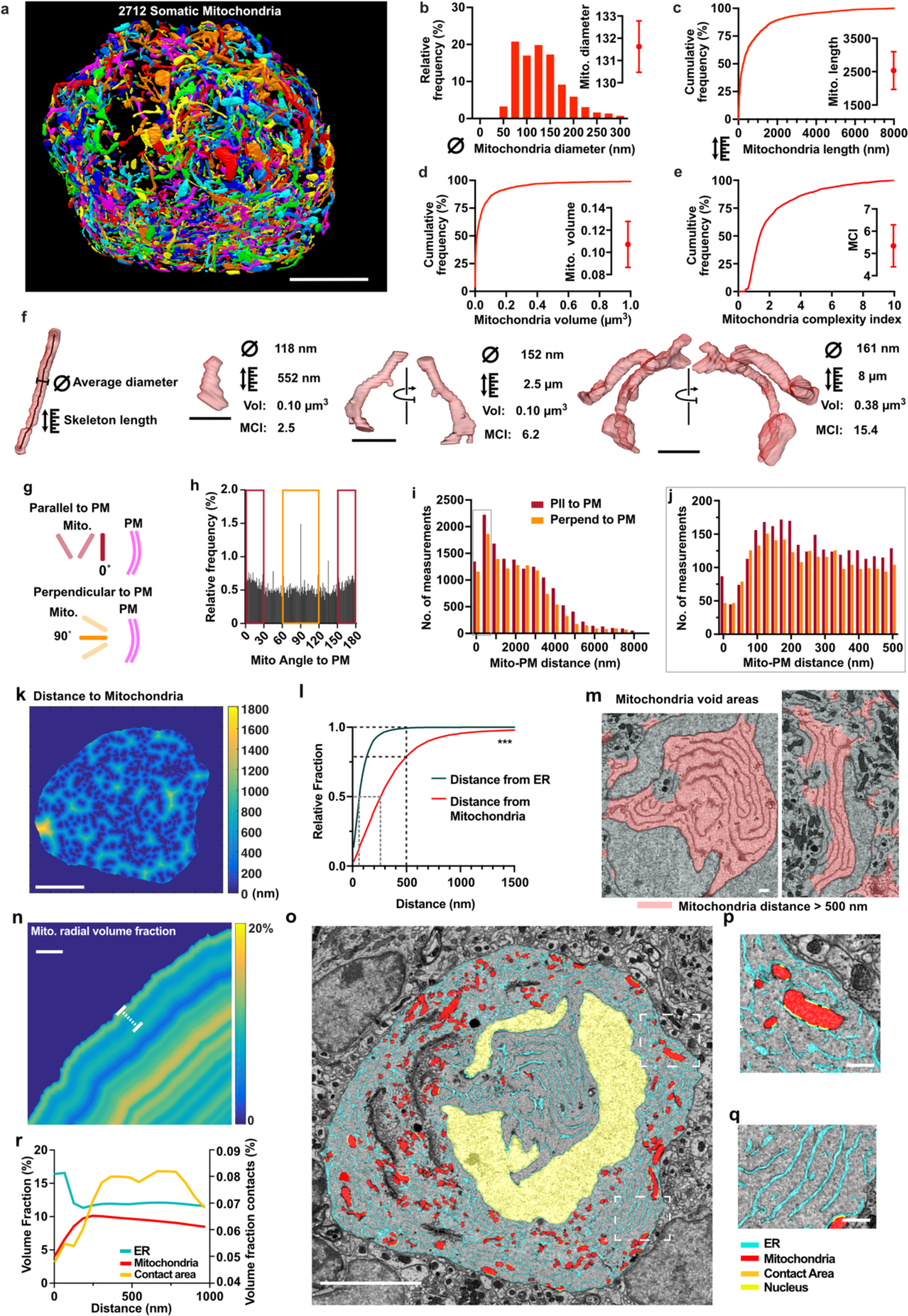
Mitochondria ultrastructure in the Soma of Purkinje neurons. (**a**) 3D color-coded rendering of 2712 mitochondria inside a single Purkinje neuron soma. (**b**) Diameter per mitochondrion (nm). (**c**) Cumulative distribution and average length with SEM of mitochondria. (**d**) Cumulative distribution and average volume with SEM per mitochondrion in µm^3^. (**e**) Cumulative distribution and average with SEM of the Mitochondria Complexity Index (MCI). (**f**) 3D renderings of representative examples for different types of mitochondria. A skeletonization version is used to determine the length and the mean diameter is determined as the closest distance to the surface at each point of the skeleton. Orientation of mitochondria was quantified relative to the PM and (**g-j**) defined as parallel (pll; red) or perpendicular (perpend; orange) to the PM. (**k**) Heatmap plot of mitochondria distance maps, shows where intracellular areas are void of mitochondria (light colors). (**l**) The cumulative mitochondria distance plot across the entire soma of the SBEM reconstruction reveals that mitochondria are less distributed than the ER throughout the soma, (**m**) leaving areas of large ER sheets without mitochondria. (**n**) Mito distance maps (**o-q**) example images (**r**) and cumulative radial distribution plot (across the entire soma) reveal that mitochondria are inwards shifted as compared to the ER, with ER-mitochondria contact sites also increasing until 250nm distance from the PM. Scale Bars: (**a,k,o**) 5µm; (**n**) 1µm; (f, p,q) 0.5µm

**Table 1.**
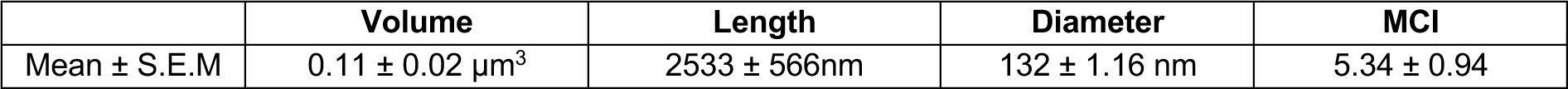
Average shape descriptor of somatic mitochondria in the Purkinje neurons.

|  | Volume | Length | Diameter | MCI |
| --- | --- | --- | --- | --- |
| Mean ± S.E.M | 0.11 ± 0.02 µm<sup>3</sup> | 2533 ± 566nm | 132 ± 1.16 nm | 5.34 ± 0.94 |

Unlike for the ER, only about one quarter of the soma is very close to the next mitochondria within 250 nm (28.03% within 250nm), meanwhile 78.33% of the soma is within 500nm of the next mitochondrion (**Figure 4 k-l**). This means that in the Purkinje neuron soma the ER is more dispersed than mitochondria throughout the soma, with effectively every region of the soma containing ER. Thus, compared to the ER, there are somatic areas comparatively far from mitochondria (**Supplementary Figure 3**). Further, we find that areas devoid of mitochondria are often where large stacks of large multi-layered ER sheets are located (**Figure 4m**).

We also characterized the radial intracellular distribution of mitochondria, by measuring their relative volume occupancy across the soma with respect to their distance to the PM (**Figure 4n-q**). We find that in contrast to the ER, mitochondria were mostly present at distances above 200nm and more from the PM (**Figure 4r**), and that only few mitochondria are in contact with the somatic plasma membrane. The ER is frequently closely associated with mitochondria, forming regions known generally as ER membrane contact sites (MCSs). It is known that these contact sites of the ER, mitochondria and cell membranes are critical for maintaining the calcium in balance in the soma and facilitate the calcium uptake into the ER and mitochondria (*8*, *33*), which influence mitochondrial function and energy metabolism and are vital for various cellular processes. To better understand the structural organization of ER–mitochondria contact sites, we quantified their radial distribution within the soma, defining contact sites as regions where the ER–mitochondria distance is less than 30 nm. We found a prominent peak in the radial distribution at 250 nm distance from the PM, which is at the intersection point of the radial peaks of ER and mitochondria (**Figure 4r**).

### The mitochondrial and endoplasmic reticulum endoarchitecture from the axon initial segment to the first heminode of Ranvier

The axon initial segment (AIS) is a thin, unmyelinated region of the axon that originates from the axon hillock and has a critical role in the neuronal physiology, as it generates action potentials in Purkinje neurons (*34*) and the length axon initial segment was shown to provide an additional mechanism for adjusting neuronal excitability (*35*, *36*). Even though several structural features have been described in the AIS, such as a discrete axoplasmic reticulum (*3*), a rather unique dense granular material coating the interior of the axon membrane (*37*) and disc-like ER structures resembling the spine apparatus (*3*, *38*), which are thought to be increased at proximity to GABAergic projections, our understanding of the overall ultrastructural organization in this zone still remains rudimentary. Since both the ER and mitochondria are storing and releasing intracellular calcium, they play a critical role in this zone for the generation of action potentials (reviewed in (*39*)). To examine the ultrastructure of the AIS in more detail we segmented the cytoplasm (**Figure 5a**), the ER and the mitochondria. We separated the AIS into sub-regions, from the internodal region (**1**), the first heminode of Ranvier with the juxtaparanode (**2**) and hemi-paranode (**3**), the distal (**4**), medial (**5**) and proximal AIS (**6**) and the axon hillock (**7**; **Figure 5b**). Along these subregions we find significant variations in the content of ER and mitochondria (**Figure 5c**). Among those the myelinated axon areas (**1&2**) have a significantly increased content in mitochondria, whereas mitochondria appear almost completely absent from the paranodal region (**3**) and the proximal AIS (**6**) (see **Figure 5c-d, g-h**). The ER content is however significantly elevated in the narrow paranode and distal AIS, while it is also decreased in the proximal AIS (**Figure 5c-d, e-f, h).** The reduced amount of both, mitochondria and ER in the proximal AIS might be related to the occurrence of the prominently visible microtubule bands in this segment (**Figure 5h**).

**Figure 5.**
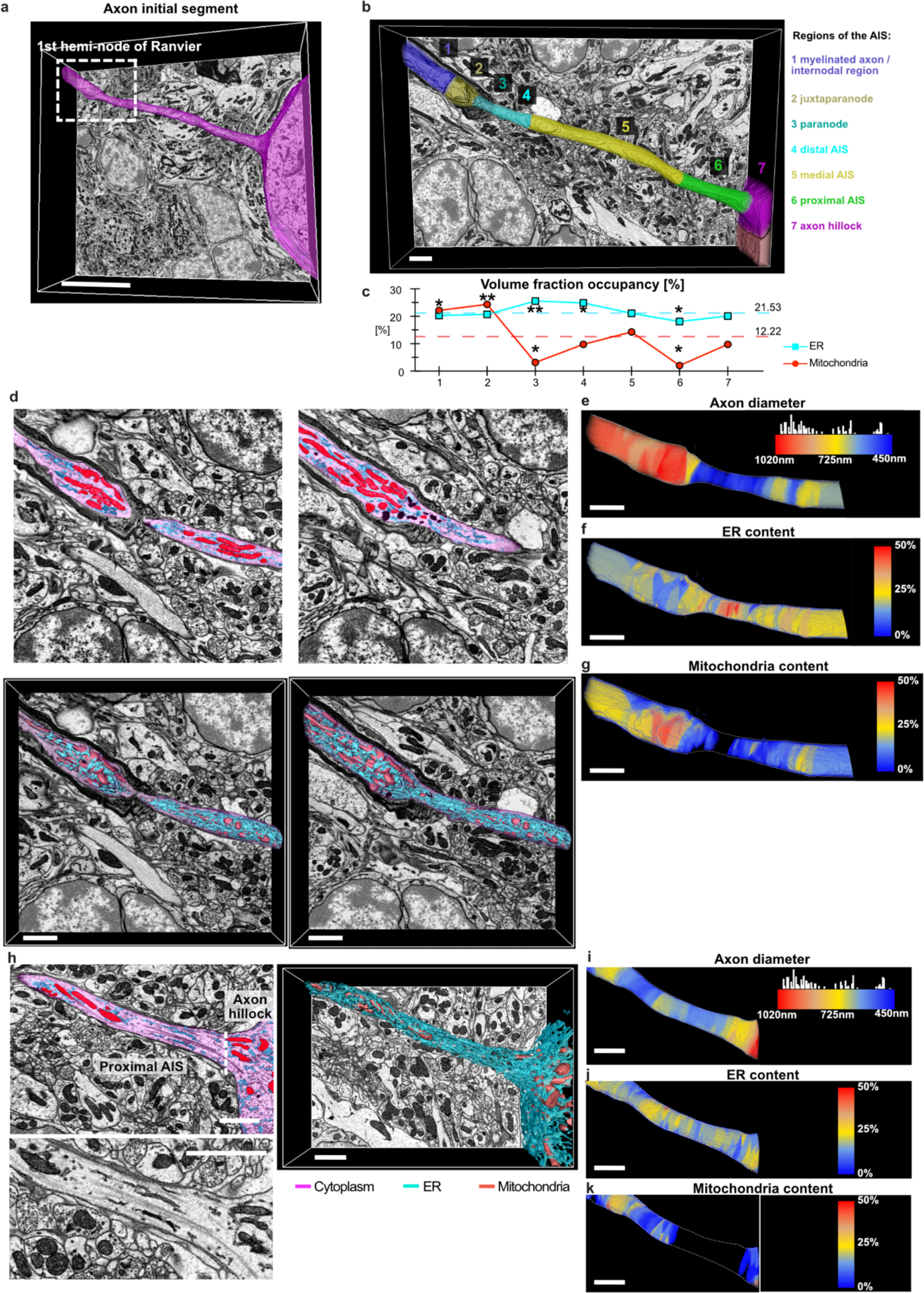
Intracellular distribution of ER and Mitochondria along the axon initial segment from the axon hillock until the first node of Ranvier. **(a**) 3D surface reconstruction of the Axon initial segment (AIS) from a SBEM volume. (**b**) The AIS was further divided into sub-regions to analyze if intracellular differences occur along the AIS. (**c**) Volume occupancy of ER and mitochondria along the subregions of the AIS shows strong variations, with absence of mitochondria in the proximal AIS (6) and close to the heminode (3). The mean contents are represented by dash lines. (**d**) 2D segmentations of sections of the hemi-node and 3D volume rendering shows the transition points at the heminode, with high ER, low mitochondria content. (**e**) Color-coded rendering of the axon diameter shows a narrowing of the axon to 450nm at the hemi-node, with a strong widening to 1µm once the axon is myelinated. (**f**) Color-coded rendering of the ER content reveals a nanodomain of very high ER-content (up to 50%) at the narrow constriction of the heminode. (**g**) Color-coded rendering of the ER content reveals that in the same nanodomain mitochondria are completely absent. Whereas the juxta-paranode contains elevated levels of mitochondria, which are more than twice the average in the AIS. (**h**) 2D section of the axon hillock and the proximal AIS shows the large microtubule filaments contained in this area. 3D rendering of the ER and mitochondria segmentations shows mostly ER and less mitochondria in the axon towards the hillock. Color-coded rendering (**i**) of the axon diameter displays a broadening of the axon towards the hillock with (**j**) continuous content of ER but (**k**) an absence of mitochondria in the proximal AIS. *: p<0.05, **: p<0.01 Scale Bars: A: 5µm. B and D-K: 1µm.

The first heminode of Ranvier has a critical role for the action potential initiation and for burst firing (*40*). Interestingly, we identify some notable differences at the anatomical level in the axon diameter of Purkinje neurons, which decreases at the heminode, from a mean diameter of 622nm (at the medial AIS) at the unmyelinated axon to 475nm at the paranode before the myelination begins. However, as soon as the myelination begins (at the juxtaparanode) the axon diameter increases to 1000nm (**Figure 5e**) and then continues with a mean diameter of 855nm (**Table 2**).

**Table 2.** Anatomy of the axon initial segment from the axon hillock until the first node of Ranvier.

| AIS region | 1<br>Internode<br>(myelinated) | 2<br>Juxta-<br>paranode | 3<br>Hemi-<br>(para)node | 4<br>Distal<br>AIS | 5<br>Medial<br>AIS | 6<br>Proximal<br>AIS | 7<br>Axon<br>hillock | Sum: | Mean: |
| --- | --- | --- | --- | --- | --- | --- | --- | --- | --- |
| Length [μm] | 1.45 | 0.99 | 0.89 | 1.81 | 6.65 | 2.94 | 2.58 | 17.31 | n.a. |
| Volume [μm<sup>3</sup>] | 2.37 | 1.43 | 0.34 | 0.74 | 4.25 | 1.95 | 3.22 | 16.68 | n.a. |
| ∅ Diameter [nm] | 857 | 855 | 475 | 534 | 622 | 644 | 505 | n.a. | 603 |

Since the ER is continuously very prominent throughout the entire heminodal area, we examined its relative content in more detail, and found it increases punctually in a microdomain at the nodal constriction, making up to 50% of the entire cytosol (**Figure 5f**), whereas it decreases towards the myelinated AIS at the perinode and the juxtaparanode back to below 25%. Conversely, the mitochondrial content decreases, and falls to a complete absence at the nodal constriction (**Figure 5g**), whereas moving along further into the juxtaparanode, once the myelination enveloped the axon the mitochondria content increases substantially to 50% and remains elevated at about 25% at the internode region, inside the myelinated axon. We further noted several degradative catabolic organelles in the juxtaparanodal and paranodal region of the AIS.

## Discussion

To date our understanding of the intracellular organization of neurons is shaped by long-standing analysis performed on 2D electron microscopy and resolution-limited fluorescence-based 3D light microscopy of selectively stained or fluorescent protein marked molecules or structures. As new techniques now allow imaging of large 3D extents with high resolution volumetric electron microscopy, a major challenge has remained for one to perform the segmentation of these data at the required scale and precision, allowing accurate quantification of details of these distinct intracellular structures and their associations with one another. The second challenge to improve our understanding the vast complexity of the intracellular organization lays in defining features to analyse, which can distil the organization principles out of a seemingly chaotic cytoplasm. As future studies will increasingly aim to understand the *endoarchitecture* within neurons, to know which are organizational principles and how they are physiologically contributing, it is first and foremost important to identify features which can be used to describe this architecture. With this study, we intend to lay out an important path for future studies to examine the organization principles of 3D neuronal ultrastructure in health and disease. Looking ahead, a complete *neuronal endoarchitectome* of adult nervous tissue will require to encompass a broad range of constituents including mitochondria, the ER, the Golgi apparatus, peroxisomes, and lysosomes. Traditionally viewed as purely catabolic organelles, lysosomes are becoming recognized as active participants in regulating anabolic processes, particularly through their interaction with the ER where their positioning was shown to directly influence ER remodeling and tubule dynamics in response to metabolic states (*41*). This highlights a bidirectional communication between degradation pathways and biosynthetic compartments. Incorporating such catabolic-anabolic crosstalk will be important for future studies aiming to understand neurodegenerative diseases. Importantly, since neurons are long-lived and do not divide, it is critical to perform these quantifications of spatially distinct patterns of the neuronal endoarchitectome in adult brain tissue as it has been shown that dynamic reorganizations such as mitochondrial mobility are much reduced in this system as compared to cultured cells or primary neuronal cultures (*42*, *43*).

When considering the neuronal energy metabolism, we discovered several interesting features in the distribution of mitochondria. For one, we found several areas of the neuron where mitochondria are absent, and therefore the energy must be supplied e.g. by diffusion of ATP from adjacent areas. While it is common knowledge that mitochondria are absent inside the nucleus, we found that, in addition, mitochondria are often absent from the areas surrounding the nucleus where large ER sheets are present, as well as from certain axonal subregions. Most noteworthy is an absence of mitochondria at the heminodal constriction, the most distal non-myelinated area of the AIS, since this has been identified to be the an area critical for the initiation of both, simple and complex spikes in Purkinje neurons (*44*). Na^+^ channels at the distal AIS are essential for the generation of action potentials (*45*) and recent studies found that the sodium channels in this area are underlying burst firing (*40*) and that their content in this area regulates the threshold for action potential generation (*46*). After an action potential however, the electrogenic Na^+^/K^+^-ATPase has to be highly active to elevate the membrane potential and to restore the ion gradient again. The absence of mitochondria of the neuron in this area is therefore puzzling, as this process is entirely ATP dependent.

Noteworthy is however also the drastic reduction in the axon diameter that we observed, which implies that a substantially smaller number of Na^+^-ions are required due to the much smaller volume (since a 30% reduction in diameter from the average axonal diameter of 0.6µm to 0.45µm already leads to a >50% reduction in axonal volume). It is interesting that in the same axonal nanodomain the ER composes 50% of the axoplasmic volume. We therefore postulate that elevated Ca^2+^ concentrations can occur in this particular nanodomain and that excluding mitochondria from this area prevents exposing them to toxic Ca^2+^ levels which would be a death signal for mitochondria (*47*). At the same time the reduced axonal volume also lowers the amount of ATP required to restore the membrane potential and excitability, making it more energy-efficient overall.

Conversely, several regions within the cell exhibit an extremely elevated mitochondrial content. Notably one of the areas with the highest mitochondria content is the juxtaparanode where up to 50% of the axoplasm is occupied by mitochondria, which provides a likely source of ATP to support the Na^+^/K^+^-ATPase activity, which is essential for sustaining to enable repeated action potentials at the first heminode of Ranvier.

## Methods

All animal procedures were approved by the Institutional Animal Care and Use Committee (IACUC) of University of California, San Diego and complied with all relevant ethical regulations for animal research.

### Tissue preparation for serial block-face scanning electron microscopy (SBEM) and electron tomography

Samples for SBEM were prepared as described in (*27*). Male mice and rats (age 4–6 weeks) were anesthetized using a mix of ketamine (Pfizer, 10mg/kg, i.p.) and xylazine (Lloyd, 2mg/kg, i.p.). They were then perfused transcardially first with Ringer’s solution at 37°C containing heparin (10 units/ml) for 3 minutes and then perfused for 15 min with a fixation solution containing 2% paraformaldehyde, 2.5% glutaraldehyde and 2 mM CaCl2 in 0.15 M sodium cacodylate buffer. Fixation was initiated at 37 °C and then cooled to 4 °C. The brain was then placed in ice-cold fixative for 2-5 h. Brain sections of the cerebellum were cut parasagittal into 100μm thick slices using a vibratome (Leica). The slices were stained with reduced Osmium (2% OsO4/1.5% potassium ferrocyanide) in ice-cold 0.15M cacodylate buffer for 1 h. After washing 3x for 5 min, the slices were placed for 20min in a filtered 1% thiocarbohydrazide (TCH) solution. The slices were rinsed 3x with ddH2O and then placed in 2% OsO4 for 30 min. The sections were rinsed with ddH2O and left in 2% uranyl acetate aqueous solution overnight at 4°C. On the following day, sections were washed 5x 2 min in ddH2O at room temperature. Slices were gradually dehydrated using a series of ice-cold ethanol solutions (50%, 70%, 90%, 100%, 100%) for 5 min each. Tissue sections were placed for 10min in ice-cold 100% acetone and 10min in 100% acetone at room temperature. Sections were infiltrated using 50% acetone–50% Durcupan ACM epoxy resin (Electron Microscopy Sciences) mix for 6 hrs. Slices were placed 3x in fresh 100% Durcupan every 6 hrs over the next day. The stained slices were embedded in 100% Durcupan with ACLAR and two mold-release coated glass slides and resin was hardened in vacuum at 60°C for 48 h.

Samples for ET were prepared as described as described in (*27*).

### Image Acquisition

#### SBEM acquisition

SBEM was performed using a Merlin SEM (Zeiss) with a Gatan 3View system at high vacuum. The SBEM volume from the cerebellum of a P40 male mouse was acquired with dimensions of 57.17 × 76.22 × 17 μm^3^. The mouse SBEM volume was acquired at 4.76nm x/y pixel spacing and 70nm section thickness. A SBEM volume of 189 × 118 × 60 µm^3^ from a rat sample (5288071) was acquired with 2.6 keV at 5.9nm x/y pixel spacing (1210x Magnification) and 40nm section thickness with a raster size of 32.000 x 20.000 pixels.

#### Electron Tomography

Five multi-tilt electron tomograms of semi-thin sections were acquired in the mouse cerebellum, covering each 24.7 - 377.3 μm^3^, from the axonal as well as the dendritic pole of Purkinje neurons at 2.1 nm - 3 nm voxel size with 0.5 degree and reconstructed at 4.2 nm - 6 nm voxel size in x/y/z and reconstructed, with following specific parameters. Tomogram 1 volume dimensions were 12.9 x 12.9 x 1.06 µm^3^ and the volume was reconstructed at 4.3 nm voxel size in x/y/z. Tomogram 2 volume dimensions were 12 x 12 x 2.62 µm^3^ and the volume was reconstructed at 6 nm voxel size in x/y/z. Tomogram 3 volume dimensions were 12 x 12 x 0.44 µm^3^ and the volume was reconstructed at 6 nm voxel size in x/y/z. Tomogram 4 volume dimensions were 8.4 x 8.4 x 0.35 µm^3^ and the volume was reconstructed at 4.2 nm voxel size in x/y/z. Tomogram 5 volume dimensions were 8.4 x 8.4 x 0.69 µm^3^ and the volume was reconstructed at 4.2 nm voxel size in x/y/z.

### Data analysis

#### General workflow and software used

Image segmentation for all cellular organelles consisted of a workflow of initial manual training data generation. Manual image segmentations and manual proof editing was performed using VAST Lite (*48*) or IMOD (*49*). Subsequently CDeep3M (*27*), a software package to train and apply a deep neural network for large-scale image segmentation, was trained on different cellular constituents. Data was computationally segmented and manually proof-edited. Subsequent data analysis and quantifications were performed in Matlab and in Python. Data rendering was performed in Amira 3D (2023.1.1). Statistical analysis and plotting of graphs were performed in GraphPad Prism. Figures were assembled in Affinity Designer (1.10.8).

#### Volume occupancies

Volume occupancies of subcellular organelles were computed per image plane by quantifying the number of segmented pixels for each structure relative to the total soma volume. The fractional occupancy of mitochondria, endoplasmic reticulum, Golgi apparatus, and nucleus was expressed as a percentage of soma area in each plane, and converted to physical volume using known voxel dimensions, with cytoplasmic volume inferred by subtraction.

#### Morphometric Analysis of the endoplasmic reticulum

Endoplasmic reticulum (ER) morphology was quantified on a per-plane basis using 2D segmentations. Each ER structure within a plane was identified as a distinct connected region and analyzed individually. For each ER strand, a skeleton representation was computed to identify its central axis. To assess the relative orientation of the ER with respect to the plasma membrane (PM), the angle of the central axis of each ER strand was calculated, along with the angle of the PM curvature at the point closest to the ER. The ER strand length was calculated by summing the pixels along its skeleton and converting the result into micrometers using the pixel dimension. To estimate the diameter, a 2D distance transform was applied to the ER segmentation, and local radii were sampled along the entire skeleton path; the mean diameter was calculated as the average of doubled distance transform values at each skeleton point. To assess the proximity of the cytoplasm to the next ER (or mitochondria) a distance map was computed from the inverse image within the soma mask. Occurring distance values were concatenated across the entire cell for the ER and for mitochondria to identify the aggregated distance within the cell to the most proximate ER or mitochondrion.

#### Mitochondrial 3D Morphometry Analysis

To assess the 3D morphology of individual mitochondria in SBEM (serial block-face electron microscopy) datasets, segmented binary volumes of mitochondria were analyzed using voxel-based and mesh-based geometric descriptors. First isolated mitochondria were isotrapically resampled and 3D skeletonization was applied to each mitochondrial volume to extract its medial axis. The total mitochondrial length was then estimated by summing the number of skeleton voxels and converting to micrometers using the voxel dimensions. To estimate the diameter, a 3D distance transform was applied to the mitochondrial volume, and local radii were sampled along the entire skeleton path; the mean diameter was calculated as the average of doubled distance transform values at each skeleton voxels. Mitochondrial volume was quantified using two approaches: voxel-based volume (via simple voxel counting) and mesh-based volume (via triangulated isosurface calculation).

The Mitochondria complexity index was calculated as described earlier (*50*) as metric to distinguish morphologically complex from simple mitochondria using following formula: 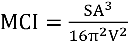 where SA is surface area and V is volume.

#### Plasma Membrane Coverage Analysis

In the electron tomography volumes, we quantified the coverage of the plasma membrane (PM) by the endoplasmic reticulum (ER). To this end we performed a 2D plane-wise analysis on binarized segmentation masks of the soma and ER. The PM was defined as the perimeter of the soma mask, extracted using morphological edge detection. To avoid misclassification at the edge of the tomogram, pixels close to the image borders were excluded from the PM analysis. For each PM pixel, the Euclidean distance to the nearest ER-positive pixel was computed using a distance transform. A PM pixel was considered “covered” by ER if it was located within 30 nm of any ER signal, based on the in-plane pixel resolution provided for the dataset. The percentage of covered PM pixels was calculated for each plane, and results were aggregated across the full image stack.

#### 3D axon diameter map

To color code the axon segmentation for the axon diameter we developed following strategy: The image stack with the binary axon segmentation was isotropically re-sampled to 36nm (or 18nm) voxelsize, and a skeletonization as well as the inverse directed 3D distance transform were calculated from the isotropic image stack. The diameter at each point of the binary skeleton is twice the 3D distance transform. Therefore, the skeleton was multiplied with 2x the 3D distance transform, multiplied by the resampled voxelsize (36nm), which results in a 3D skeleton encoding the axon diameter at each centerpoint of the axon. To revert the diameter of the axon to its 3D extent we calculated the index map of the closest point on the skeleton for the entire axon. Each point of the axon was then reassigned with the value of the closest point on the 3D skeleton encoding the axon diameter. The result is a 3D axon diameter map, which was subsequently volume rendered in Amira (displayed in **Figure 5**).

#### 3D axonal mitochondria and ER content maps

As outlined above for the axon diameter map, we also encoded the content of mitochondria and ER. To this end we calculated in a similar manner for each voxel of the axon the closest voxel on the 3D skeleton, and the percent of the axonal voxels belonging (being closest) to each skeleton voxel containing mitochondria or ER respectively. The resulting maps encode the content of mitochondria or ER along the axon (shown in **Figure 5**).

#### 3D axonal volume fraction content

The volume fractions across the different regions of the AIS were calculated by dividing voxels that are mitochondria or ER in the region by the total voxels in this region. One sample t test was used to compare segments to the population mean using GraphPad Prism.

## Authors’ contributions

MGH and MHE conceived and guided the work. MGH, SP, TD and EB performed experiments. MGH performed computational segmentations. MGH, SVS, EPC, KR, YW, LM, BLB performed manual proof-editing of computational segmented data. MGH wrote code to perform computational analysis of the data, did the meshing, rendering and assembly of the figures and drafted the manuscript. All authors read, edited and approved the final manuscript.

## Acknowledgements

This work was supported by grants from the NIH’s NIGMS (R01GM082949) and NINDS/BRAIN Initiative (U24NS120055) and the NSF (2014862) to MHE. This work was supported by a Visible Molecular Cell Consortium (VMCC) fellowship to MGH. This study was further supported by the German Research Foundation (Deutsche Forschungsgemeinschaft; DFG) project ID 509099330 to MGH and SVS and funding from Germany’s Excellence Strategy-Exc-2049-390688087 and the DZNE e.V. to SVS.

---

**Supplementary Figure 1.**
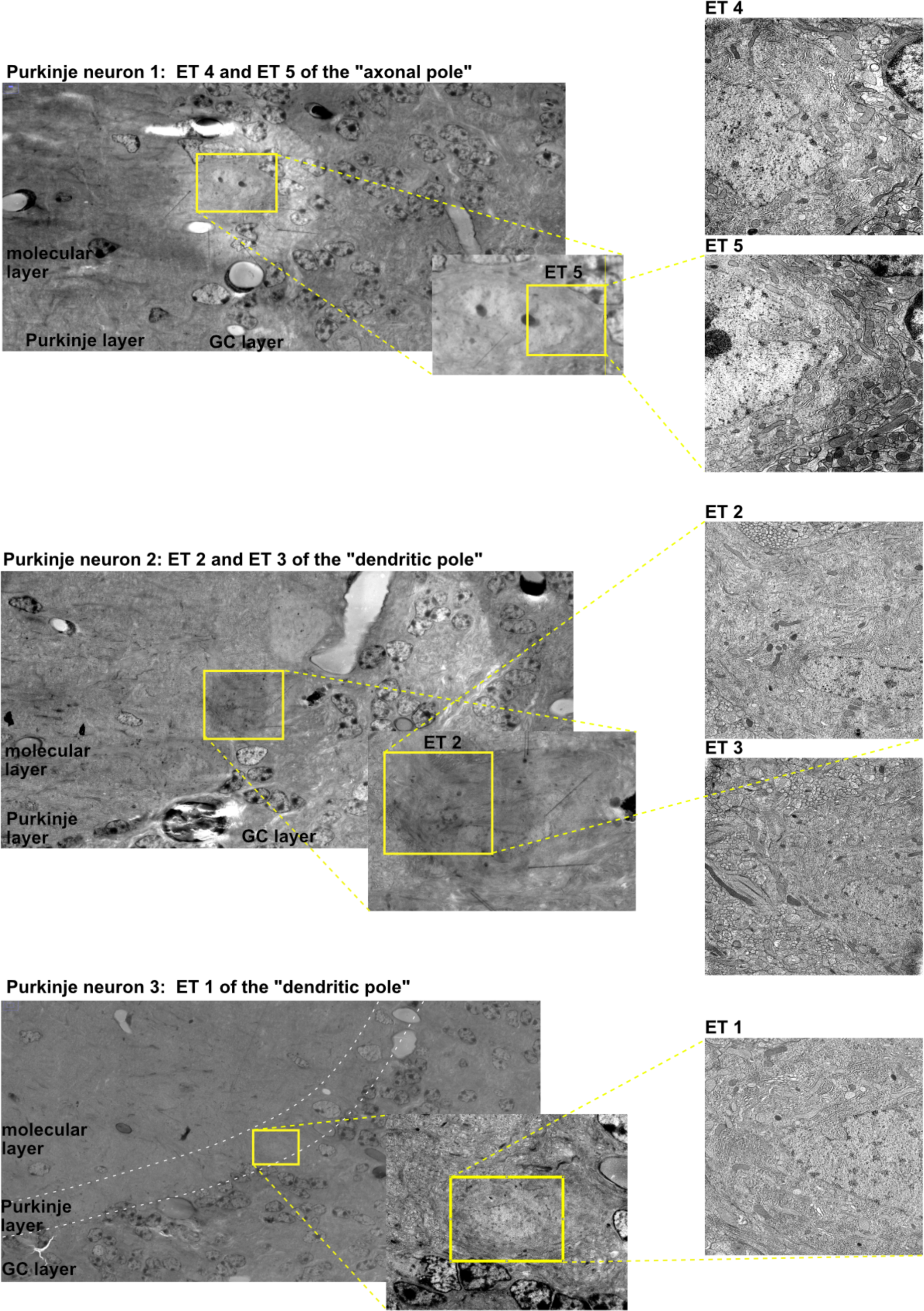
Cerebellar region overview for electron tomography volumes. Electron tomography volumes were acquired from the dendritic or axonal pole of different Purkinje neurons.

**Supplementary Figure 2.**
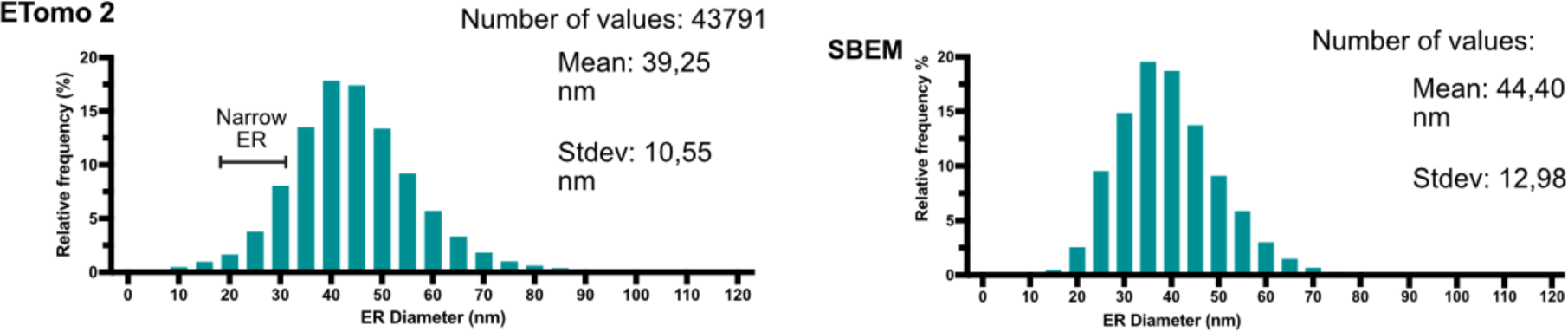
ER diameter quantification using high-pressure frozen freeze substituted electron tomography and chemically fixed SBEM.

**Supplementary Figure 3.**
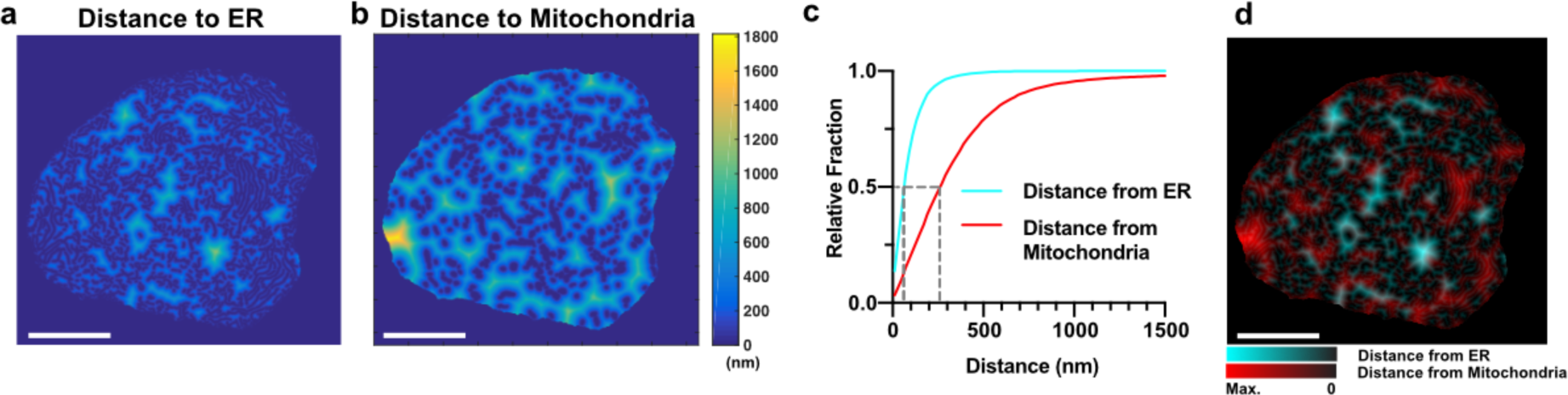
Comparative examination of ER and Mitochondria void areas in the Purkinje neuron soma. To identify the proximity of the entire soma to the closest mitochondrion and the closest ER we calculated a closest proximity distance map of (a) ER and (b) Mitochondria. (c) The cumulative distance from the closest ER and mitochondria reveals a significantly closer proximity to the ER than to the Mitochondria (Kolmogorov-Smirnov test; p<0.0001), where 50% of the soma has ER within 60 nm distance, and 50% of the soma has mitochondrion within 250 nm distance. (d) Overlaying the distance maps of both showed that there is little to no overlap between the areas that are very distant from ER and the areas that are very distant from Mitochondria. Scale Bars: 5μm

